# In situ characterization of the 3D microanatomy of the pancreas and pancreatic cancer at single cell resolution

**DOI:** 10.1101/2020.12.08.416909

**Authors:** Ashley Kiemen, Alicia M. Braxton, Mia P. Grahn, Kyu Sang Han, Jaanvi Mahesh Babu, Rebecca Reichel, Falone Amoa, Seung-Mo Hong, Toby C. Cornish, Elizabeth D. Thompson, Laura D. Wood, Ralph H. Hruban, Pei-Hsun Wu, Denis Wirtz

## Abstract

Pancreatic ductal adenocarcinoma (PDAC) is one of the deadliest forms of cancer. Accumulating evidence indicates the tumor microenvironment is highly associated with tumorigenesis through regulation of cellular physiology, signaling systems, and gene expression profiles of cancer cells. Yet the mechanisms by which the microenvironment evolves from normal pancreas architecture to precursor lesions and invasive cancer is poorly understood. Obtaining high-content and high-resolution information from a complex tumor microenvironment in large volumetric landscapes represents a key challenge in the field of cancer biology. To address this challenge, we established a novel method to reconstruct three-dimensional (3D) centimeter-scale tissues containing billions of cells from serially sectioned histological samples, utilizing deep learning approaches to recognize eight distinct tissue subtypes from hematoxylin and eosin stained sections at micrometer and single-cell resolution. Using samples from a range of normal, precancerous, and invasive pancreatic cancer tissue, we map in 3D modes of cancer invasion in the tumor microenvironment, and emphasize the need for further 3D quantification of biological systems.

## Introduction

The growth and spread of invasive cancer, and the relationship of invasive cancers to pre-existing structures such as vessels, nerves, and ducts, is best understood through accurate three dimensional (3D) representations.(1-3) Pancreatic ductal adenocarcinoma (PDAC) is one of the deadliest forms of cancer, with a 5-year survival rate of only 9%.(4) PDAC arises from well-characterized precursor lesions in the pancreatic ducts, is surrounded by an immunosuppressive desmoplastic stroma, and has a proclivity for vascular invasion and metastasis to the liver.(5) These biological factors are insufficiently understood when studied in two dimensions (2D), as it is difficult if not impossible to infer information such as connectivity and 3D cell density and morphology from 2D media. Therefore, increased knowledge of the 3D microenvironment of the pancreas and changes to this microenvironment with progressive tumorigenesis will lead to a better understanding of the underlying biology of pancreatic cancer. While many surrogates for studying tumorigenesis in 3D have been developed in vitro and in vivo, (6-10) quantitative 3D study of naturally occurring cancers in human tissues (cancer in situ) is generally lacking. Data derived from analyses of cancer within large 3D human tissue samples, in particular, have the potential to significantly increase our knowledge of the human tumor microenvironment.

Recent advances in tissue clearing techniques have been employed to explore human diseases in 3D.(11-16) In pancreatic cancer, tissue clearing has been used to show that sustained epithelial-to-mesenchymal transition is not required for vascular invasion.(2) However, inconsistent clearing and poor antibody penetration into the dense desmoplastic stroma that characterizes PDAC, as well as limits on the size of tissues that can be successfully cleared, hinder the power of tissue clearing techniques when applied to pancreatic cancer.(13, 16) The reconstruction of serial, hematoxylin and eosin (H&E) stained sections has also been implemented to study disease in 3D.(17-19) Though time-consuming manual annotations or costly immunohistochemical (IHC) labeling or mass spectrometry has been introduced to identify biological components in 3D,(17, 18) the costs and time associated with labelling cellular and structural bodies in serial sections largely limits its applicability.

Here, we demonstrate a novel method for the 3D reconstruction and quantification of large tissue volumes at single-cell resolution, which we name CODA. CODA digitally reconstructs 3D tissues from scanned, serially sectioned H&E tissue sections using image registration techniques. The method incorporates deep learning semantic segmentation to label eight distinct tissue types of the human pancreas without incorporation of additional stains. CODA is also capable of single-cell analysis and is utilized here to quantify in 3D cellular content and spatial distributions amongst different non-neoplastic and neoplastic tissue types, information that is important in the design of early detection tests.(20) With CODA, we analyzed normal pancreas tissue and pancreas tissue containing cancer precursors and PDAC in tissues of cm-dimension and μm-resolution. We derived detailed 3D insight of PDAC development and progression in situ.

## Results

### CODA: 3D reconstruction and labelling of serial histological sections

To study pancreatic cancer invasion, we identified a human pancreas sample (S04-PDAC) containing poorly differentiated infiltrating ductal adenocarcinoma immediately adjacent to a large region of grossly normal pancreas (Supplementary Table 1). The formalin-fixed paraffin-embedded sample was serially sectioned every 4µm. Sections were stained with H&E and digitized at 20x magnification, providing x and y (lateral) resolution of 0.5µm and z (axial) resolution of 4μm (Figure 1A). We developed CODA, a workflow for the digital reconstruction of these serial tissue images into a high-resolution, multi-labelled tissue volume amenable to the extraction of high-content cellular and non-cellular information.

**Figure 1.**
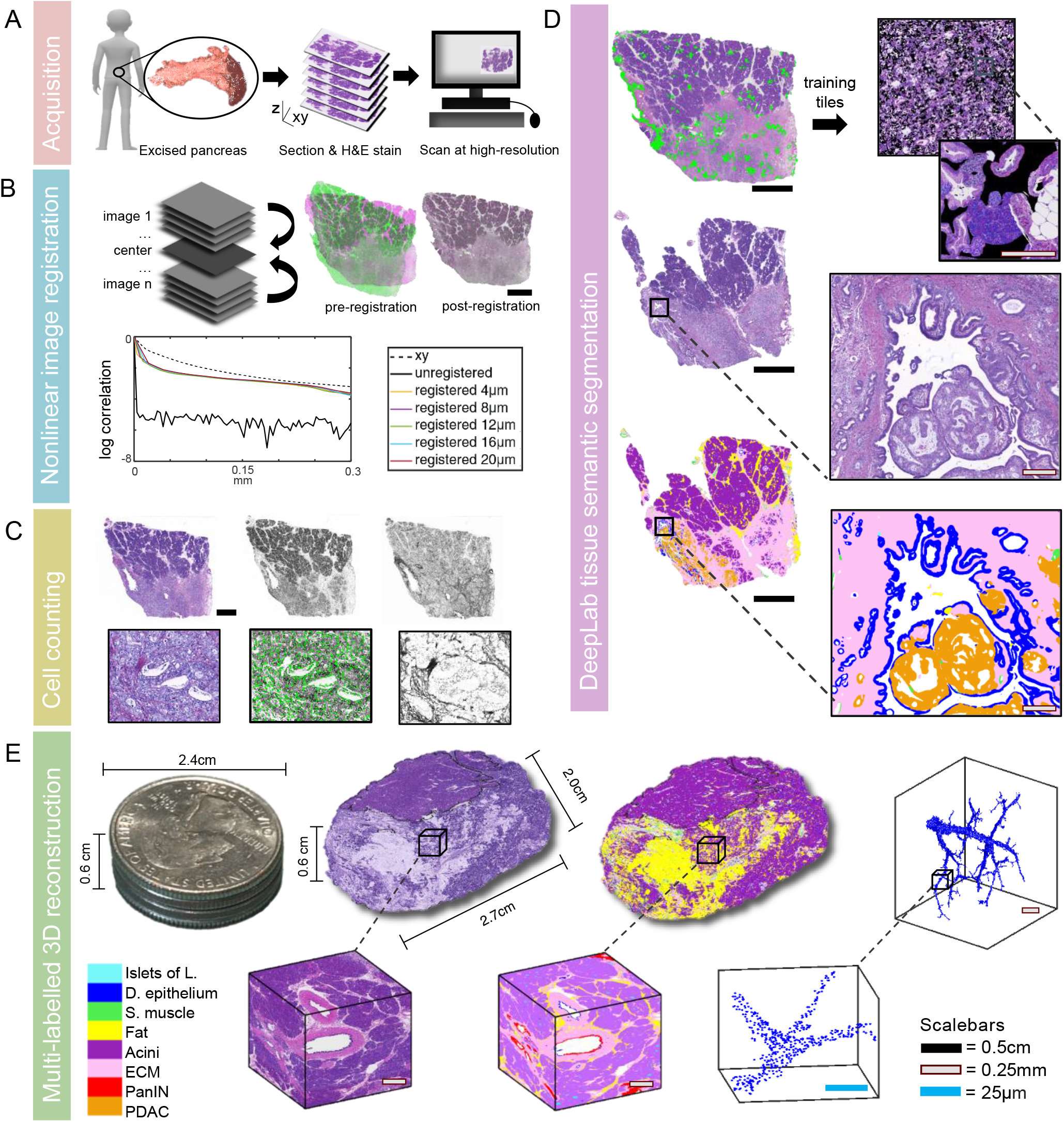
CODA. A Human pancreatic tissue is formalin fixed, paraffin embedded (FFPE), serially-sectioned, stained, and scanned at high resolution. B Tissue images are registered to create a digital volume. Correlation of tissue image intensity in the xy dimension of single tissue sections is used as a reference for registration quality. Correlation of intensity in the z dimension in unregistered image stacks, and registered image stacks with different z-resolutions show that we maintain 99% registration quality with a z resolution of 12µm. C Cells are identified using the hematoxylin channel of color deconvolved images. 2D serial cell counts are corrected using the in-situ measured nuclear diameter of cells in different tissue bodies. D DeepLab deep learning semantic segmentation is created using manual annotations of tissue types which are randomly overlaid on large black tiles for training. Tissue images are then labelled to a resolution of 2µm. E 3D reconstruction of >1000 serially sectioned pancreas tissue. 3D renderings are created at the cm, mm, and µm scale at both the tissue and single cell level.

First, the independent serial images were mapped to a common coordinate system using image registration. Tissue sections were coarsely aligned using whole field affine registration. However, reconstruction of serial histological sections is complicated by the malleability of tissue, which unpredictably stretches, folds, and splits to produce nonuniform deformation between z-planes.(21) Therefore, we further applied an elastic registration approach to account for local tissue warping. Elastic registration approaches compute nonlinear transformations and have been successfully used to register histological images.(17-19, 22) Uniquely, our approach was developed and optimized for the registration of pancreas tissue sections and incorporates downsampling to increase its speed. We found the registration process performs similarly between consecutive tissue sections or tissue sections up to five z-planes apart (Figure 1B, Supplementary Figure 1), allowing us to improve the throughput of the workflow by processing only one in three serial images. Overall, the registration workflow serially aligned the S04-PDAC tissue sample containing 1,499 serial histological sections in 3 h.

We further established an automated cell detection workflow to locate all nuclei in each histological section based on color deconvolution and a previously established algorithm described in ref.(23) During validation, automated cell detection performed similarly (94% consistency) to manual cell annotation (Figure 1C, Supplementary Figure 2A). In situ diameters of each cell type were measured and incorporated to extrapolate true 3D cell counts from cell counts on serial 2D z-planes (Supplementary Figure 2B).

To visualize and quantify the architecture of the pancreas, labeling distinct tissue subtypes in the volume is crucial. Deep learning methods have been successfully used to identify many structures in H&E images, such as inflammation,(24) cancer cells,(25, 26) and extracellular matrix (ECM).(27) We established a deep learning workflow utilizing DeepLab semantic segmentation and a pretrained ResNet50 network (Supplementary Figure 3A) (28) to label eight pancreas tissue subtypes recognizable by trained pathologists in H&E images without additional molecular probes. A total of eight tissue subtypes were identified: normal ductal epithelium, precursors (pancreatic intraepithelial neoplasia [PanIN] or intraductal papillary mucinous neoplasm [IPMN]), PDAC, smooth muscle & nerves, acini, fat, ECM, and islets of Langerhans (Figure 1D). Importantly, to increase model accuracy, we create our training dataset by semi-randomly overlaying extracted annotated regions on a large image, then cutting this large image into many training and validation images, allowing us to control the heterogeneity of class appearance in the dataset (Suppelentary Figure 3B). The trained deep learning model achieved precision and recall of >90% for each class in S04-PDAC on independent testing images (Supplementary Figure 4) and labelled the serial sections at a resolution of 2µm/pixel in 36 h. Registration of the detected cell coordinates and labelled images allowed us to create cm-scale multi-labelled tissue volumes with μm and single cell resolution which could be easily visualized and analyzed quantitatively.

### 3D assessment of pancreatic cancer invasion in situ

The fully labeled reconstructed volume and detected cells allowed us to assess the dimensions of the sample, composition of tissue subtypes, and number of cells in each subtype quantitatively (Figure 2A). The S04-PDAC sample had estimated dimensions of 2.7cm x 2.0cm x 0.6cm with a total volume of ∼2.2 cm^3^. The sample contained ∼1.1 billion cells. Of these, 2.1 million cells (∼0.2%) were cancer precursor and ∼10.5 million cells (∼1%) were invasive cancer.

**Figure 2.**
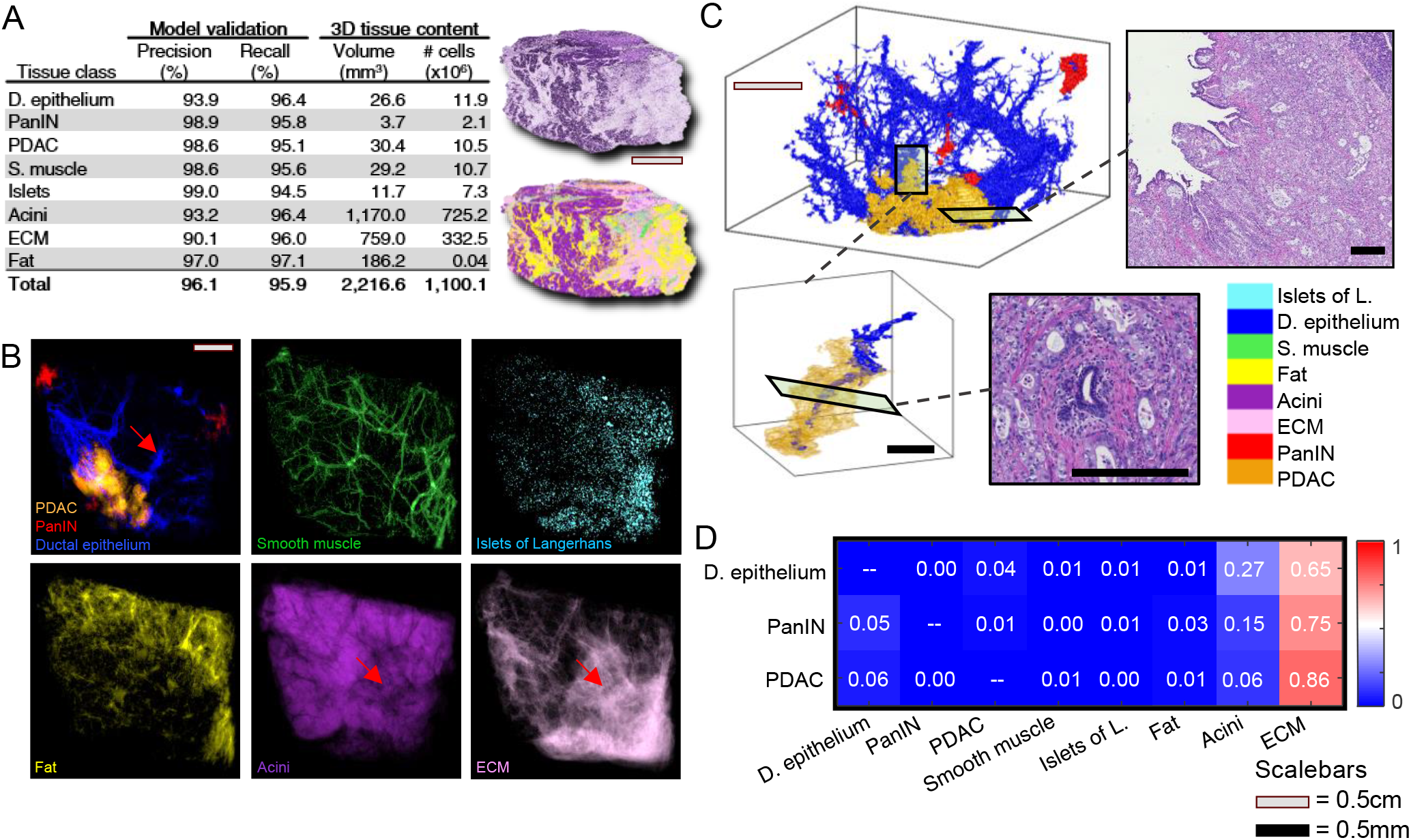
3D reconstruction of cancerous human pancreas allows quantitation of cancerization of large duct. A Deep learning training accuracy was assessed using manual annotations of tissue subtypes and model was iteratively trained until subtype precision and recall of >=90% were obtained. Bulk tissue subtype volume and cell counts were calculated. B z-projections of classified regions show a normal pancreatic duct extending from a large cancer mass to an area of acinar atrophy (red arrows). C 3D reconstruction and sample histology show cancerization of a large duct and a cancer protrusion growing along a smaller duct. D Quantification of tissues present in 50µm surrounding ducts, precancers, and cancer show ECM surrounds all three tissue subtypes and increases in quantity with progression from normal duct to precancer to cancer.

We then visualized the landscape of cancer invasion at the leading edge of the cancer and adjacent normal tissue via z-projections and 3D renderings. The z-projections of the normal duct and benign spindle shaped cells (vasculature and nerves) showed well-connected tubular morphology (Figure 2B). The z-projections showed a large mass of adenocarcinoma located at one side of the tissue sample that had a strong spatial association with a large normal pancreatic duct. The 3D rendering of PDAC, precursor lesions, and normal ductal epithelium revealed that this spacial association was in part because the invasive cancer had infiltrated the ductal epithelium, a process known as cancerization of the ducts(29) (Figure 2C, Supplementary Video 1). A smaller, non-neoplastic duct fed into the portion of the duct colonized by the invasive cancer, and this upstream pancreatic parenchyma was atrophic with acinar drop-out and increased content of ECM and prominent islets of Langerhans (Figure 2B, red arrows).(30) The cancer and atrophic region identified using the deep learning 3D reconstruction was confirmed by manual review of the histology (Figure 2C), validating the 3D reconstruction and labelling capabilities of CODA.

In addition, we found three small projections of the invasive cancer extending from the larger tumor into surrounding normal pancreatic tissue, appearing to colocalize smaller pancreatic ducts. Examination of the 3D rendering revealed that the largest of these projections surrounded and extended parallel to a pancreatic duct for a distance of >3mm without invading the epithelial layer (Figure 2C, Supplementary Video 2). This observation was further confirmed by manual review of the sample histology. Overall, the visualization of the leading edge of cancer in a large 3D pancreas sample indicates that invasive cancers can track in the periductal stroma parallel to pre-existing ducts in the pancreas, as has been hypothesized in 2D.(31, 32)

### Interpatient analysis for exploration of cancer tumorigenesis in 3D

To further investigate 3D patterns in pancreatic cancer tumorigenisis, we characterized the changes in tissue architecture in four additional tissue samples for a total of five samples spanning S01-Normal: normal pancreas; S02-PanIN and S03-IPMN: pancreas containing precursor lesions PanIN or IPMN; S04-PDAC: pancreas containing invasive poorly differentiated pancreatic ductal adenocarcinoma with adjacent grossly normal tissue; and S05-PDAC: pancreas containing invasive poorly-differentiated pancreatic ductal adenocarcinoma with no adjacent normal tissue (Supplementary Table 1, Supplementary Videos 3 and 4). Using CODA, we obtained multi-labeled 3D maps of these tissue samples (Figure 3). Individual deep learning models were trained for each sample with performance of >90% class precision and recall compared to manual annotations (Supplementary Figure 4).

**Figure 3.**
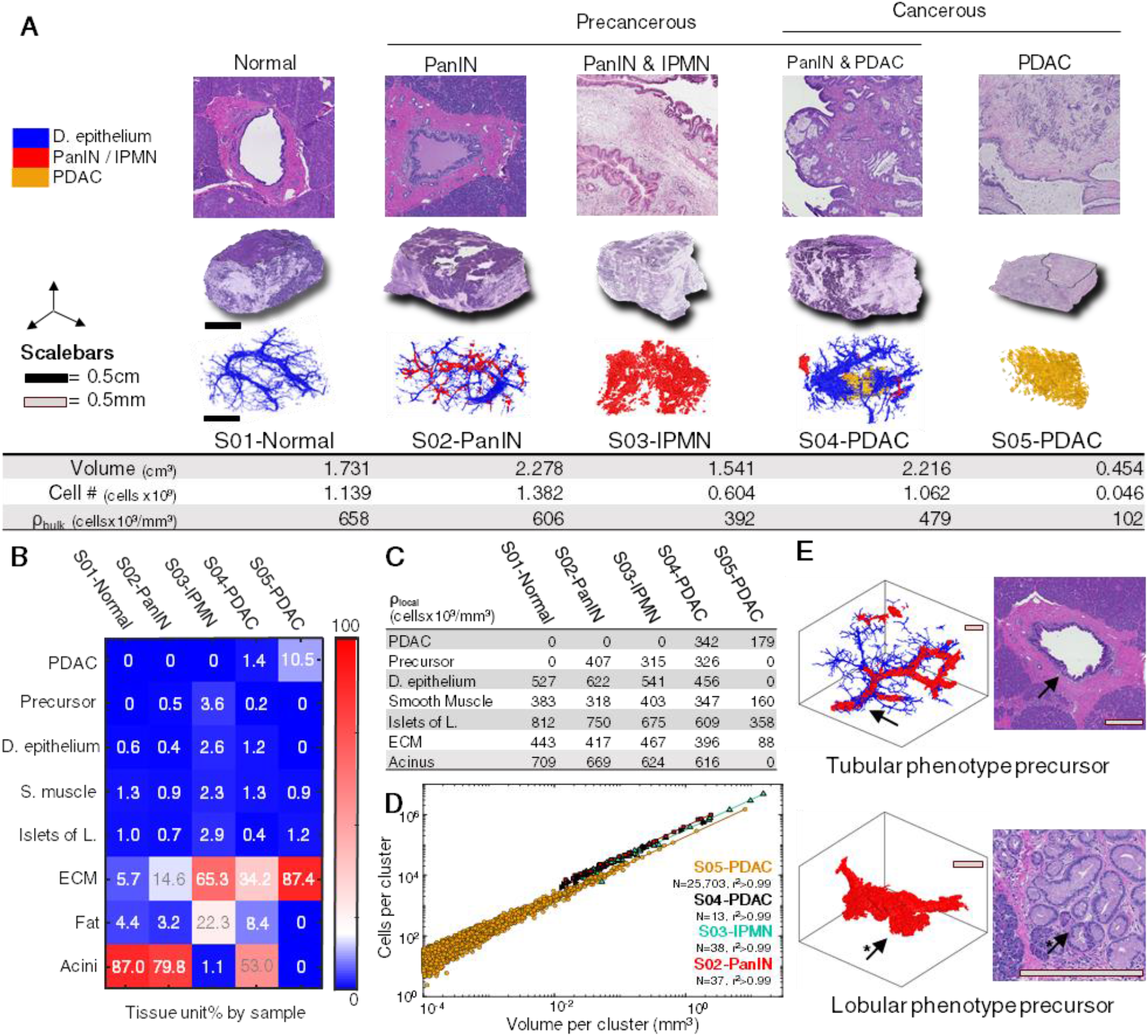
Inter-patient pancreas analysis from cm-scale to single cell resolution. **A B**ulk tissue volumes, cell counts, and cell densities for samples containing normal pancreas, precancerous lesions, and pancreatic ductal adenocarcinoma. **B** Heatmap showing tissue subtype percentages of tissue samples. **C** Table showing local tissue subtype cell densities. **D** Plot of volume per cluster vs. cells per cluster for independent precursor and cancerous cell clusters across four tissue samples. Lines of best fit show that precancer and cancer clusters maintain similar cell density independent of cluster volume. **E** 3D renderings and sample histology illustrate two 3D phenotypes of PanIN observed. Tubular PanIN preserve normal pancreatic ductal morphology, while lobular PanIN resemble clusters of acinar lobules.

Our approaches provide direct 3D visualization of normal pancreas, pancreatic cancer precursors (PanIN and IPMN), and PDAC at cm-scale with μm and single-cell resolution (Figure 3A). Through quantification of tissue volume and cell count, we were able to compare overall cell densities between samples. Counterintuitively, we found that with progression from normal pancreas to cancer precursor to cancer, bulk cell density (ρ_bulk_, total number of cells normalized by total tissue volume) decreased. Comparison of ρ_bulk_ with tissue subtype percentages revealed that tissues containing precursors and invasive cancers, which had the lowest ρ_bulk_, contained the highest percentage of ECM and lowest percentage of acini (Figure 3B). Markedly, acinar content dropped significantly from normal (87.0%) in the S04-PDAC case (53.0%), which contains cancer and adjacent grossly normal tissue, and was nearly absent in both S03-IPMN (1.1%), and S05-PDAC (0%), as the normal pancreatic parenchyma in these samples was entirely atrophic. ECM content was highest (87.4%) in the case of extensive infiltrating PDAC (S05-PDAC) compared to S01-Normal pancreas (5.7%). Therefore, although growth of precancerous and invasive cancer cells implies an increase in cellular content, cocurrent acinar atrophy and the laying down of desmoplasic stroma and connective tissue result in an overall decrease in bulk in situ cell density with development of pancreatic cancer.

In addition to bulk cell density, we utilized local cell density (ρ_local_) for deeper exploration of these structural changes (Figure 3C). We define ρ_local_ as the cell density of a tissue subtype within the detected volume of that subtype – while calculation of bulk cell density shows the number of cells in the total tissue volume, ρ_local_ allows exploration of the closeness or sparseness of tissue subtypes at a local level. We found that ρ_local_ decreased in the acini, islets of Langerhans, ECM, normal ductal epithelium, and precursor subtypes with progression from normal pancreas to PDAC, suggesting that these cells are larger or more sparse in cancer precursor and cancerous samples than they are in the normal sample. Indeed, direct visualization of the histologic slides by a pancreatic pathologist confirmed that normal ductal epithelial cells appeared larger in the S04-PDAC sample than in the normal and precancerous cases, and that normal acinar cells packed more tightly than atrophic acinar cells. Though intuitively one might expect inflammation in precancerous and cancerous tissues to result in higher ρ_local_ in the ECM, importantly, we found the opposite to be the case. While we observed local regions of dense inflammatory cells near precursor and cancer cells, cocurrent growth of less-inflammed ECM as extracellular fibrous connective tissue replaced atrophied acini, and desmoplastic stroma developed around the cancer resulted in a counterintuitive decrease in ECM cell density with tumorigenesis.

### Structural changes in human pancreas precancers with tumorigenesis

In addition to bulk measurements, we utilized CODA for enumeration of architectural patterns in the samples. Pancreatic intraepithelial neoplasia by definition involves the complex branching of the pancreatic duct system. In 2D it can be impossible to discern if one is observing two separate PanIN lesions or one PanIN that has branched, or whether a PanIN occupies a small region of a pancreatic duct or extends for many mm within the ductal architecture. Readily, we found in 3D that precursors present in a range of volumes, can be architecturally simple or highly branched, and that many spatially distinct precursors can develop within cm-scale regions. We identified 37 PanIN lesions in sample S02-PanIN, 38 precursors in S03-IPMN, and 13 PanIN lesions in S04-PDAC, varying in size from 0.013-9.7mm^3^ and containing a range of 4,000-3,728,000 cells. Surprisingly, we found precursor ρ_local_ to be relatively constant and independent of volume, with mean precursor cell density of 404,000±1,000 standard error cells/mm^3^ in sample S02-PanIN (Figure 3D). Similarly, we found cancer ρ_local_ to be independent of tumor volume, with mean cancer cell density of 189,000±300 standard error cells/mm^3^ in S05-PDAC, for cancer cell clusters containing a range of 1-1,500,000 cells. This suggests that pancreatic cancer precursor and cancerous cells occupy the same amount of space whether they present in-situ as single cells or within very large tumors.

While assessing 3D connectivity of cancer precursors, we identified two 3D structural phenotypes of PanIN which we term tubular and lobular (Figure 3E, Supplementary Video 5). Tubular PanIN lesions appeared as elongated, ductal, branching structures, while lobular PanIN lesions appeared as clumped, “bunches of grape-like,” near-solid masses. Review of the corresponding H&E sections by a pancreatic pathologist revealed that tubular PanINs resided within more proximal pancreatic ducts, while lobular PanIN lesions resided at the terminal junctions between ducts and acinar lobules. The lobules in these cases representing areas of acinar to ductal metaplasia. Upon further investigation we found that nearly a third of PanINs exhibited both phenotypes, with regions of growth within both more proximal pancreatic ducts and more distally as the ducts merged with acini. Our analysis suggests that the structural appearance of PanIN mirrors the appearance of the tissue it develops within. Tubular PanINs resemble the shape of pancreatic ducts, while lobular PanINs take on the architecture of acinar lobules. While it is known that PanIN can extend from the ductal epithelium to foci of acinar to ductal metaplasia,(33, 34) our analysis suggests that this involvement of the acinar tissue affects the 3D organization of the precursor.

These findings emphasize that the dramatic, volumetric changes to the organization of pancreas tissue brought on by development of precancers and invasive cancers are large – when analyzing changes with development of pancreatic cancer, we observed decreases to overall cell density, decreases to ρ_local_ for many tissue subtypes, increase in regions of acinar atrophy and ECM deposition, and complex 3D morphological phenomena occurring on cm-scale.

### Collagen alignment in tissue is a fundamentally 3D concept

The layer of ECM that surrounds normal pancreatic ducts is called the ductal submucosa and can be clearly observed in 3D renderings of pancreatic ductal structure in the S01-Normal sample (Figure 4A). Analysis of the tissue composition of the immediate surroundings (within 50μm) of the cancer of sample S04-PDAC showed that 85% of the surrounding tissue was ECM compared to 75% around PanIN and 65% around normal ductal epithelium (Figure 2D). This calculation, along with the identification of a 3mm growth of cancer along the outside of a normal duct (Figure 2C) showed that the progression and invasion of PDAC is strongly associated with the ECM.

**Figure 4.**
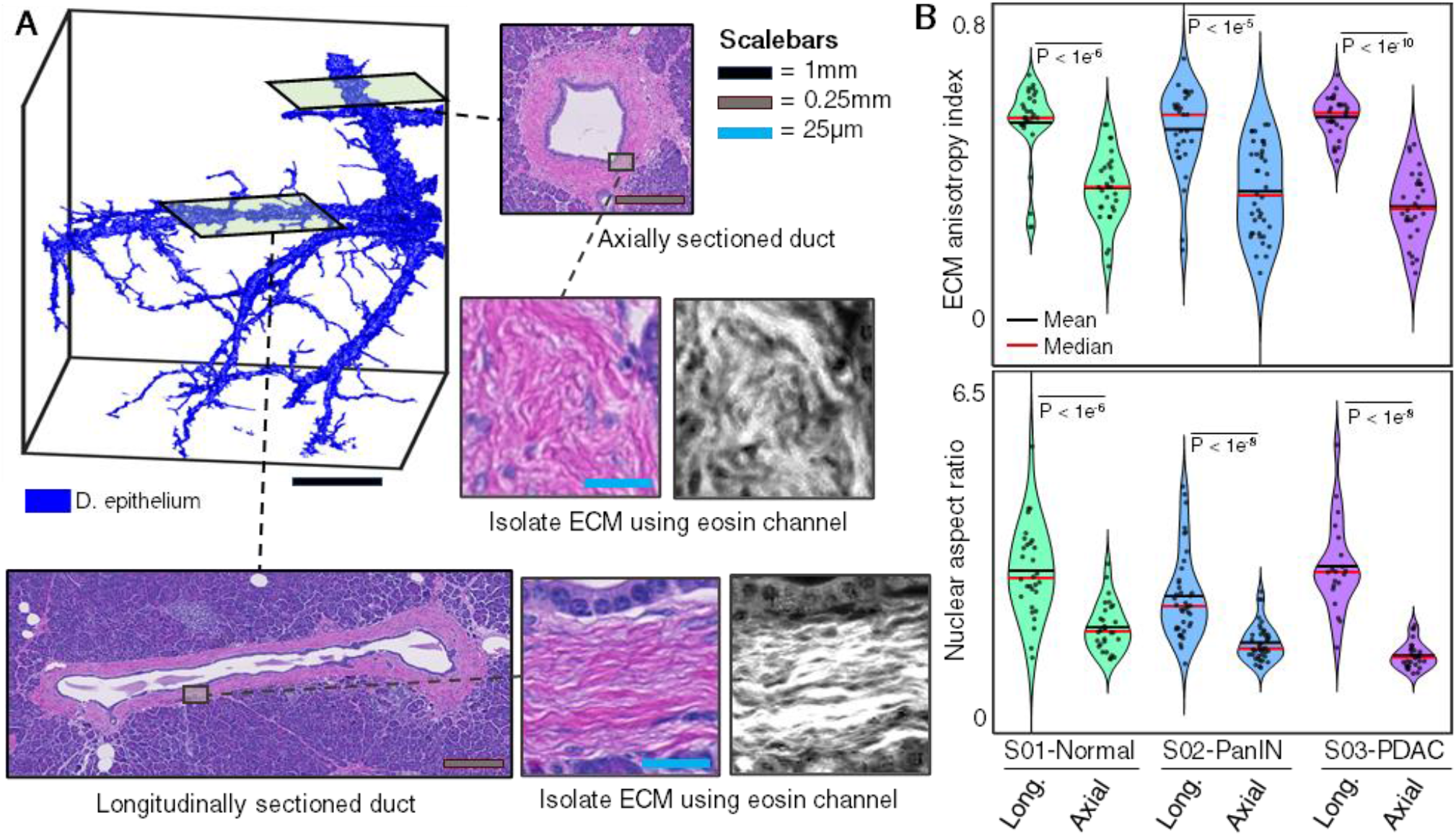
3D rendering of pancreatic ducts emphasizes 3D nature of measured collagen alignment. **A** 3D reconstruction of serially sectioned pancreatic ducts allows quantitative analysis of nuclear morphology and collagen alignment in context of the 3D organ. 18 histological sections which intersect pancreatic ducts axially or longitudinally were located, consisting of nine axial ducts and nine longitudinal ducts from three patient samples. Nuclei and collagen fibers were isolated using color deconvolution. **B** Axially and longitudinally sectioned ducts were compared to assess collagen alignment. ECM anisotropy index, representing the alignment of local collagen fibers, was significantly higher in longitudinal ducts in all three cases, suggesting collagen is measurably straighter in the longitudinally sectioned ducts. Nuclear aspect ratio was significantly higher in longitudinal ducts in all three cases, suggesting stromal cells are more elongatedin longitudinally sectioned ducts.

Previous studies have shown that ECM alignment has a role in mediating PDAC invasion, and that patients with highly aligned desmoplastic stroma surrounding pancreatic cancer have a worse prognosis than patients with poorly aligned desmoplastic stroma.(35-37) Having observed the growth of invasive pancreatic cancer parallel to non-neoplastic pancreatic ducts, we further examined apparent ECM alignment on histological sections in the ductal submucosa as a function of the orientation of the pancreatic duct in 3D. Navigating the 3D renderings of the ductal epithelium in three samples (S01-Normal, S02-PanIN, and S04-PDAC), we identified coordinates where the ductal submucosa was cut at two extremes: perpendicular to the long axis of the duct (axially-sectioned), or parallel to the long axis of the duct (longitudinally-sectioned). We located the identified 3D coordinates on the original, serially-sectioned histology images, collecting samples of a total of 18 ducts sectioned axially or longitudinally.

As expected, the periductal fibroblasts in ducts sectioned longitudinally were highly elongated in appearance compared to their round shape in axially-sectioned ducts (Figure 4A). Strikingly, we found that collagen fibers were visibly more aligned and elongated in longitudinal cross-sections than in axial cross-sections. Quantification of collagen fiber alignment using a method described in ref,(38) and calculation of nuclear aspect ratio further confirmed this observation showing significant increase in ECM alignment and nuclear aspect ratio in longitudinally sectioned ducts (Figure 4B). This finding was consistent across three tissue samples. Together, these results show that the ductal submucosa is highly aligned, like layers of an onion, along the duct’s axial direction. Additionally, the results offer an explanation for the observed pattern of cancer growing parallel to pancreatic ducts which fits the accepted model of PDAC growing along regions of aligned collagen.(32) These results highlight the importance of 3D analysis as previous studies performed in 2D have suggested there is insignificant collagen alignment around normal pancreatic ducts.(32, 36)

### 3D relationship between pancreatic cancer and blood vessels

Cancer intravasation is a known critical step in metastasis. The classical structural view of cancer intravasation is that cancer invades through the basement membrane and ECM into the vasculature.(39) However, models of the mechanism and extent of pancreatic cancer intravasation are hindered by their lack of quantification of large, 3D, in situ environments. CODA presents a direct way to visualize and quantify cancer intravasation at the cm and single-cell scale in situ.

We examined how PDAC is associated with vasculature in two large samples comprising the leading edge of poorly differentiated pancreatic cancer (S04-PDAC) and the bulk tumor region of poorly differentiated pancreatic cancer (S05-PDAC) in situ (Figure 5A). In S04-PDAC, we detected no clear patterns of cancer involvement with the vasculature, instead observing a large cancer mass with small blood vessels running throughout it (Figure 5B). We could observe no sign of cancer intravisation, and, due to the cancer’s high density in the area around the blood vessels, could not observe apparent correlations between cancer growth and blood vessel orientation. Conversely, in S05-PDAC sample, review of 2D sections by an expert pancreatic pathologist revealed an area of venous invasion (Figure 5C, Supplementary Video 6). Reconstruction of this region in 3D shows that the PDAC both surrounded and fully occluded the vein for a length of over 1.5 mm, extending out of view off both z-boundaries of the tissue sample. We quantified the density of cancer cells as a function of distance from the center of the vein, and found that cancer ρ_bulk_ within the lumen of the vessel is 6.5x the average global cancer cell density. High-resolution H&E images showed small clusters of PDAC cells distributed homogeneously within the tunica media of the vessel. Interestingly, we found eight separate instances of PDAC breaching the wall of the vein (Figure 5D), suggesting that intravasation and extravasation are not single-instance events.

**Figure 5.**
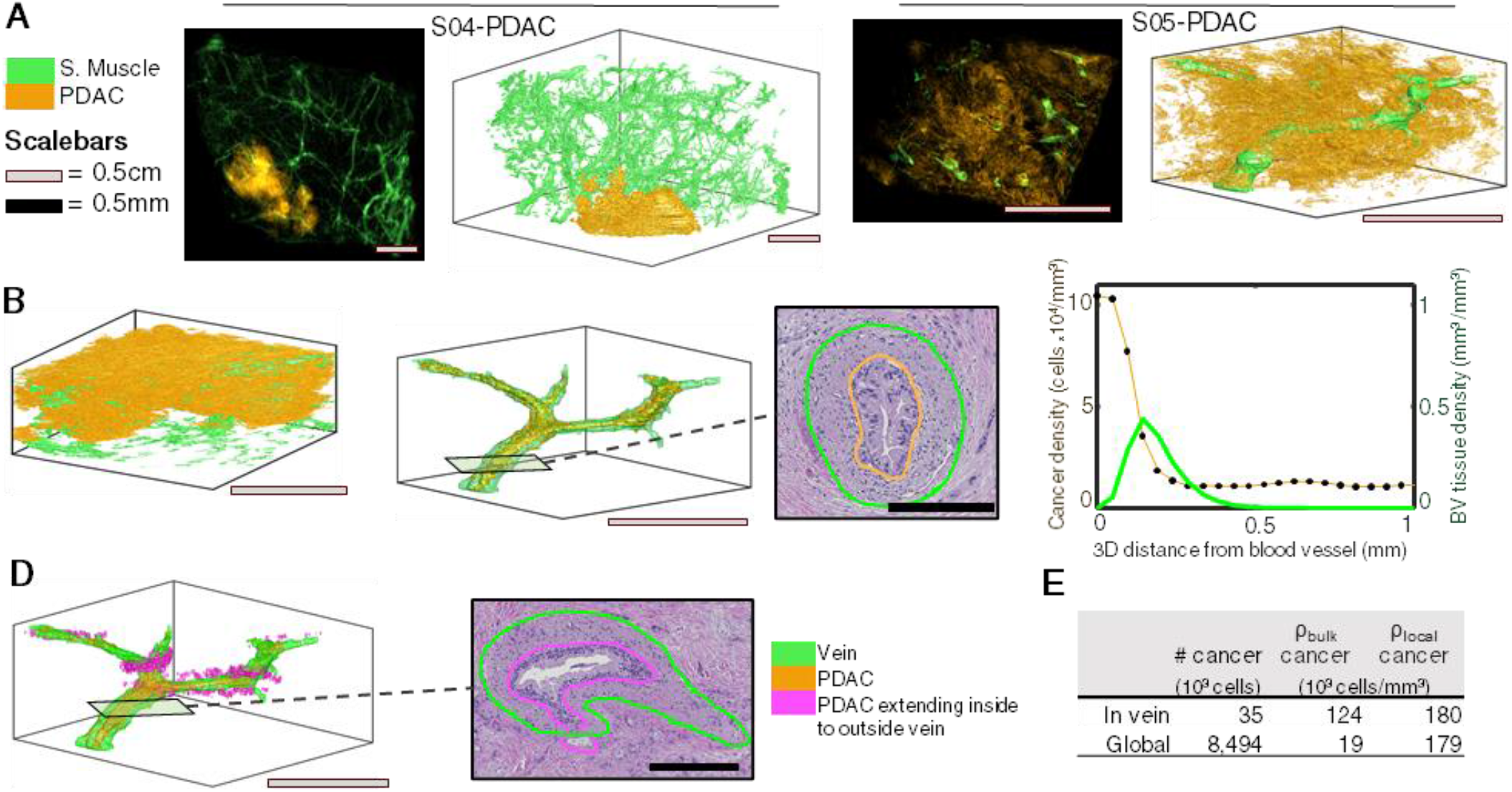
Analysis of relation of cancer to blood vessels reveals inter-patient heterogeneity. The relationship between PDAC and smooth muscle was compared between the S04-PDAC and S05-PDAC samples. **A** Z-projections and 3D reconstruction overlays of cancer and smooth muscle were created. Analysis of blood vessels in the S04-PDAC sample showed little spatial association with cancer, while a large region of venous invasion was found in S05-PDAC. **B** Small blood vessels were observed within the cancerous region of S04-PDAC, but no obvious alignment or invasion was observed. **C** Venous invasion in sample S05-PDAC was isolated, 3D reconstructed, and quantified. Cancer bulk cell density was high inside the lumen of the vein and steeply dropped off at the vessel wall and within the tissue outside the vessel. **D** Cancer detected to be breaching the vein was detected in the volume and verified at nine instances on histological sections. **E** Bulk and local cancer cell density were quantified within the vein and within the entire tissue. Bulk cell density is cancer cell count normalized by the entire volume of the vein lumen, while local cell density is cancer cell count normalized by cancer cell volume within the lumen. Bulk cell density was found to be 6.5x greater inside the vessel than the global tissue, while local cell densities inside and outside the vessel were similar, suggesting cancer cells are in closer proximity to one another inside the vessel, but that they occupy a similar volume inside and outside the vessel.

Further, we counted 35,000 cancer cells growing within the 1.5mm long region of the vein (Figure 5E). We found that while cancer ρ_bulk_ inside the vein is much higher than cancer ρ_bulk_ averaged over the entire tissue, cancer cell ρ_local_ remains constant – here, cell density is number of cancer cells normalized by total tissue volume; cancer ρ_local_ is number of cancer cells normalized by cancer cell volume. Therefore, even though cancer cells are in closer proximity to each other inside the the vein than they are in the bulk of the tissue (cell density), the cells individually take up the same amount of volume both inside and outside the vein (cell ρ_local_), i.e. the higher density of cancer cells inside the blood vessel does not force them to pack more tightly than they would outside the blood vessel. This finding stresses the importance of investigating vascular invasion in 3D, as only by comparing bulk cancer cell density to local cancer cell densityover large volumes were we able to note changes to cancer cell organization in the tissue region.

## Discussion

In this work, we show multi-labeled 3D reconstructions of human pancreas samples that provide highly-detailed structural insight of pancreatic cancer tumorogenesis within five large patient samples. CODA allows well-quantified study of in situ cancer progression at cm-scale with μm and single-cell resolution. We showed that many disconnected PDAC precursor lesions can develop within cm-scale regions of a tissue sample, and that neoplastic ρ_local_ is independent of precursor volume or 3D structural phenotype. In a single sample at the leading edge of PDAC, we found that the cancer extended furthest from the central tumor along existing, well-aligned ECM structures such as those surrounding pancreatic ducts. We emphasized the need for 3D insight in digital pathology research through quantification of collagen alignment in pancreatic ducts cut at different angles. Overall, we demonstrate that CODA provides a powerful alternative to tissue clearing for study of 3D tissue microarchitecture.

Tissue clearing is a popular approach for the study of 3D tissues, wherein intact samples are rendered semi-transparent, stained, and imaged using confocal or light-sheet microscopy.(11-16, 40) Tissue clearing techniques have been used to conduct landmark scientific research such as the imaging of all cells in a whole mouse brain(11, 12) and to assess tumor and tumor-associated macrophage heterogeneity in samples containing lung carcinoma.(41) Clearing techniques are suitable for analyses requiring few labels, as imaging of cleared tissues is often constrained to 1-5 markers per sample, with more markers feasible in µm-scale samples and fewer markers feasible in mm-scale or whole organ samples;(16, 42) and for experiments where qualitative analyses are sufficient, as inconsistent clearing and antibody penetration (especially in stiff, stromal tissues such as cancer samples, or in samples of mm or cm scale) makes quantification of imaged tissues difficult.(16, 42) For these reasons µm-scale samples and qualitative analyses are most common.(42)

Current serial sectioning methods bypass some of the shortcomings of tissue clearing methods, albeit through introduction of new challenges. Serial sectioning methods overcome the size limitations and inconsistent staining of tissue clearing by cutting tissue samples into thin (4-5µm) slices that are individually stained and scanned – there is potentially no limit to the size of tissue that can be reconstructed, and we show success here in reconstructing tissues of >2cm^3^. However, the act of cutting tissue into many thin sections introduces discontinuity to the samples, as sections can warp and fold in unpredictable ways, requiring introduction of sophisticated image registration techniques. Additionally, many serial sectioning methods rely on additional techniques for tissue labelling, including IHC staining, mass spectrometry, and manual annotation.(17, 18, 21) While these techniques contribute to the complexity and expense of serial sectioning methods, they also highlight one of its advantages: that quantification of tissues and single cells from histological sections is a popular and successful field of study in the scientific community.(24, 27, 43, 44) While groups conducting tissue clearing research often must invent new methods of analyzing complex 3D images, serial sectioning quantification can take advantage of previously developed 2D computational approaches, as serial histological samples can be quantified at the single section level and the results extrapolated to the registered digital tissue volume. Thus, while quantification of stains is simpler in current serial sectioning methods than it is in tissue clearing methods, the acquiring of tissue labels through expensive labelling methods and the necessity of sophisticated image registration techniques have hindered the general adoption of serial sectioning methods for the study of 3D tissue microarcitecture.

CODA incorporates nonlinear image registration and deep learning techniques to create multi-labelled tissue volumes using H&E images alone, avoiding the need for additional stains for tissue labelling. Here we show successful detection of eight pancreatic tissue subtypes using H&E images. By making our registered tissue dataset publically available, we leave open the possibility that future methods capable of distinguishing additional subtypes (such as immune cells or fibroblasts) from H&E sections might add additional labels to the samples analyzed here. This knowledge transfer is not possible in cleared tissue samples where unlabelled tissues cannot be visualized. Finally, our results demonstrate the ability of CODA to derive quality 3D reconstructions while skipping at least two intervening sections. Therefore, future addition of IHC labeling, gene mutation, and gene expression imaging to the intervening sections can increase the number of labels beyond what is currently discernable in H&E – the number of tissue and molecular phenotypes that CODA can label is a feat that is currently unachievable through tissue clearing or current serial sectioning approaches. A true “multi-omic” 3D map is now possible.

Overall, our analysis of cm-scale pancreas samples emphasizes the potential for 3D assessment to improve our understanding of tumorogenesis. We showed that CODA outperforms tissue clearing methods in its ability to create easily quantifiable tissue volumes, allowing quantification of deceptively simple concepts such as 3D cell count and density, vascular connectivity, tumor branching and morphology, and cm-scale tissue heterogeneity. While analogous metrics are routinely used for quantification of cell density, spatial correlation, and tumor infiltration in 2D, we show that not only are these measures different when measured in 3D, but argue that often 2D correlates are fundamentally flawed. For instance, it is impossible to accurately assess the connectivity of branching ductal structures such as PanIN and IPMN (which are distinguished by size in 2D), as we have shown that complex glandular lumina can present as distinct objects separated by centimeters of tissue on single histological sections. Indeed, the heterogeneity of the pancreatic cancer environment dictates that it is impossible for single histological sections to accurately represent the complex milieu of cancer cell growth and corresponding cell death, desmoplastic tissue development, and immune cell invasion within tumor regions. CODA is a powerful complement to previous tissue clearing methods and an upgrade to current serial sectioning 3D reconstruction methods as it designed with ease-of-quantifiability of these 3D concepts in mind.

## Materials and Methods

### Tissue acquisition and scanning

Formalin-fixed, paraffin-embedded samples were sectioned every 4μm. Every third tissue section was staine using hematoxylin and eosin (H&E), with two sections every three held out. All tissues of a single sample were scanned for validation that skipping two sections maintained registration and reconstruction accuracy. Tissues were scanned at x20 using a Hammamatsu Nanozoomer.

### Image registration

Cases contained series of tissue images scanned at 20x, corresponding to approximately 0.5μm/pixel. Openslide software was used to save reduced size copies of each image, corresponding to 8μm/pixel using nearest neighbor interpolation.(45) For each sample, the center image was identified as the point of reference (image_n_), and global and elastic registration was calculated for all other images in the sample.

We performed registration on greyscale, Gaussian-filtered, down sampled (80μm/pixel resolution) versions of the high-resolution histological sections. Global registration transformations for a pair of preprocessed tissue images was found through iterative calculation of registration angle and translation via maximization of cross-correlation. Radon transforms of the images taken at discrete angles between 0 and 359 degrees were calculated. The maximum of the cross correlation of radon transforms of the images yielded registration angle, and the maximum of the cross correlation of the rotated tissue images yields translation. Elastic registration was obtained by calculating rigid registration of cropped image tiles at 1.5-mm intervals across the globally registered images at 8μm/pixel resolution. The resulting local, rigid registration fields were interpolated and smoothed to produce a nonlinear, elastic registration transformation.

Rigid global registration was performed to sequentially register each image_n+/-m_ to the three next closest images to center, image_n+/-(m+1)_ image_n+/-(m+2)_, and image_n+/-(m+3)_. Quality of each of the three global registrations was assessed by comparing pixel-to-pixel correlation between the moving and each reference image. The registration with the best result was kept and the other two discarded. Following global registration, elastic registration was employed between the moving image and chosen reference image to create a nonlinear displacement map. This process was repeated for all images in a sample such that all images were elastically registered to the coordinate system of the center image_n_.

### Identification of cells in histological samples

First, the hemotoxylin channel of all H&E images was extracted using color deconvolution. Openslide software was used to save reduced size copies of all tissue images, corresponding to 2μm/pixel using nearest neighbor interpolation. For each image, the tissue region of the image was identified by finding regions of the image with low green channel intensity and high red-green-blue (rgb) standard deviation. Next, rgb channels were converted to optical density. Using kmeans clustering analysis, 100 clusters were identified to represent the optical densities of the image. The most common blue-favored optical density was chosen to represent the hemotoxylin channel, and the most common red-favored optical density was chosen to represent the eosin channel. The background optical density was fixed as the inverse of the average of the hemotoxylin and eosin optical densities. These three optical densities were used to deconvolve the rgb image in to hemotoxylin, eosin, and background channel images. Using methods described in ref,(23) the hemotoxylin channel images were smoothed, and 2D intensity minima of a designated size and distance from each other were identified as nuclei.

A total of 3 2mmx2mm regions were extracted from each case for validation. For each region, cells were manually located using an annotation function built in MATLAB 2020b. A manually identified cell was considered to be equivalent to an automatically detected cell if the coordinates were within 4μm of each other (corresponding to 3 pixels in the 2μm/pixel downsampled images used for cell detection). This validation showed a 94% consistency between manually and automatically detected cell coordinates.

### Deep learning tissue multi-labelling

A deep learning model was created for each case using manual tissue annotations of that sample. Openslide software was used to save reduced size copies of all tissue images, corresponding to 2μm/pixel using nearest neighbor interpolation. Seven tissue images equally spaced within each sample were extracted. For each of the seven images, we manually annotated 50 examples of each identified tissue subtype using Aperio ImageScope, creating .xml files of annotation coordinates. Annotation coordinates were loaded into MATLAB 2020a using publicly available software and were downsampled to correctly overlay on the 2μm/pixel tissue images.(46)

In order to reduce the heterogeneity of the H&E images, the H&E stain of all tissue images in each case were normalized. Using the hemotoxylin and eosin channel images created for the cell counting analysis and the optical density calculated for a reference H&E image from the same case, we reconstructed rgb images of each tissue type to the same optical density. Incorporation of image color normalization allowed us to avoid catastrophic failure of the semantic segmentation on unannotated images with drastically different staining patterns.

Bounding boxes of all annotations were identified and each annotated rgb image region was extracted and saved as a separate image file. A matrix was used to keep track of which bounding box images contained with annotation tissue types. Training images were built through creation of a 9000×9000×3, zero-value rgb image tile. Annotation bounding boxes containing the least represented deep learning class were randomly overlaid on the blank image tile until the tile was >65% full of annotations and such that the number of pixels of each deep learning class was approximately even. Annotation bounding boxes were randomly augmented via rotation, scaling by a random factor between 0.8-1.2, and hue augmentation by a factor of 0.8-1.2 in each rgb color channel. The 9000×9000×3 image tile was then cut into 324 500×500×images. 20 such large images were built, half with augmentation, to create 6480 training images, and 5 additional images were built to create 1620 validation images. 324 testing images were created using manual annotations from an image not used for training or validation.

Following dataset creation, a resnet50 network was adapted for DeepLab v3+ semantic segmentation (28) and trained to a validation patience of 5. If 90% tissue subtype precision and recall was not obtained, additional manual annotations were added to the training images and the process was repeated until desired accuracy was reached. Once a satisfactory deep learning model was trained, all tissue images in the sample were semantically segmented to create labelled tissue images with a pixel resolution of 2μm/pixel.

### 3D reconstruction of samples

Multi-labelled images created by the DeepLab portion of the CODA pipeline were consolidated into a 3D matrix using the H&E image registration results. Similarly, cellular coordinates counted on the unregistered histological sections were consolidated into a 3D cell matrix using the H&E image registration results. 3D renderings of the labelled tissue regions were visualized using the patch and isosurface commands in MATLAB 2020b and using a color scheme with a unique rgb triplet for each tissue subtype. Dimensions of rendered tissues were calculated in xy using the pixel resolution of the original x20 scanned histological sections (approximately 0.5μm/pixel) and using the tissue section spacing (4μm) in z. The resolution of the 3D renderings was 2μm/pixel in xy, the resolution used for image semantic segmentation, and 12μm/pixel in z, as only one in three tissue sections were used in the analysis. Single cells were visualized within the 3D renderings using the scatter3 command in MATLAB 2020b. For all calculations performed on the 3D labelled matrices of the tissues, the 3D matrix was subsampled using nearest neighbor interpolation from original voxel dimensions of 2×2×12μm^3^/voxel to an isotropic 12×12×12μm^3^/voxel.

### Construction of z-projections

The 3D labelled matrices of each patient case were used to construct z-projections of each tissue subtype. For each tissue subtype, the pixels of the 3D matrix corresponding to that subtype were summed in the z-dimension, creating a projection of the volume on the xy axis. The projections were normalized by their maximum and visualized using the imagesc command in MATLAB 2020b using the same color scheme created for visualization of the 3D tissue.

### Calculation of tissue spatial associations

Spatial associations of different tissue subtypes were calculated using these 3D matrices. First, a 3D matrix containing the tissue subtype of interest was isolated. Next, the regions containing that tissue subtype were dilated to a distance of 48μm. Spatial association of that tissue subtype to other tissues in the case were calculated as the percentage of each tissue subtype present in the dilated region divided by the total volume of the dilated region (not including any portion of the dilation that extended outside the tissue volume).

### Calculation of tissue content, bulk cell density, and local cell density

Tissue content was calculated by counting the total number of voxels in the isotropic 3D matrix corresponding to each tissue subtype and dividing those numbers by the total number of voxels in the tissue region of the 3D matrix. Cell density of each tissue subtype was calculated by combining the tissue subtype data in the multi labelled 3D matrix with cell coordinate data in the cell 3D matrix. Cells at each voxel in the cell 3D matrix corresponded to the tissue subtype label in the multi labelled 3D matrix (for example, a cell is labelled an epithelial cell if the nuclear coordinate was identified in a region labelled as epithelium by the deep learning pipeline). Measurements of nuclear diameter were used to estimate true 3D cell counts from the 2D cell coordinates. Using Aperio ImageScope, 100 nuclei of each tissue subtype were measured for each case. The estimated 3D cell count (C_3D_) of cells counted on serial histological sections analyzed every 3 sectionswas calculated using the formula:

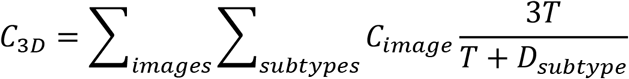

where C_image_ is the cell count for a given tissue image, T is the thickness of the histological section, and D_subtype_ is the measured diameter of a nucleas for a tissue subtype. For each tissue subtype, bulk 3D cell density was calculated by dividing the 3D extrapolated cell count of a particular subtype divided by the total volume of the tissue. Local 3D cell density was calculated by dividing the 3D extrapolated cell count of a particular subtype divided by the volume of that particular tissue subtype.

### Determination of tissue connectivity and of spatially distinct precursor lesions

The 3D multi labelled matrices were used to determine tissue connectivity. Following classification, all objects labelled as pancreatic precancers lesions or pancreatic cancer were visually verified to be precancers by inspection of the histology. Independent precursors were identified in the 3D multi labelled matrix using the bwlabeln command in MATLAB 2020b. bwlabeln identifies and labels spatially distinct objects in matrices. We calculated connectivity using bwlabeln on both the precancers alone and the precancers plus the normal ductal epithelium. Distinct precancers and cancers identified using bwlabeln could then be quantitatively analyzed or 3D rendered independently from other precancers.

### Calculation of collagen fiber alignment and fibroblast aspect ratio

Using the 3D renderings of the pancreatic ductal epithelium, we identified six regions comprising three axially and three longitudinally sectioned regions of the ducts in three cases. We located the 2D histological sections using 3D coordinates of the identified regions and cropped the region of interest from the corresponding 20x H&E images. We applied the color deconvolution method described above to the cropped 20x H&E image to separate the hematoxylin and eosin channels. We measured the alignment index of the eosin channel to compare the degree of collagen alignment in axially and longitudinally sectioned regions of the ducts. Alignment index is measured using the method described in ref.(38) An alignment index of one represents completely aligned matrix of fibers and an alignment index of zero represents an isotropic matrix of fibers. We measured the alignment index at 10 locations of each cropped image. We found each location contained an average of approximately eight cells. We manually measured the length of major and minor axis of nuclei in the ductal submucosa to calculate aspect ratios using ImageJ. In total, we measured 1546 nuclei. Violin plots were constructed from data using code available in ref. (47)

### Calculation of 3D radial density

3D radial density of tissue subtypes and cells was calculated using the multi labelled and cell coordinate 3D matrices. First, a region of interest was identified in the 3D multi labelled matrix. A logical 3D matrix was created containing only this region. Next, dilations of a predefined step size (such as 12μm) were performed. For each dilation, the number of cells and percent of each tissue subtype present in the dilation were calculated and normalized by the total volume of the dilation. A scatter plot was created with normalized tissue subtype or cell density on the y axis and distance from the region of interest on the x axis.

## Supplementary Tables

**Supplementary Table 1.**
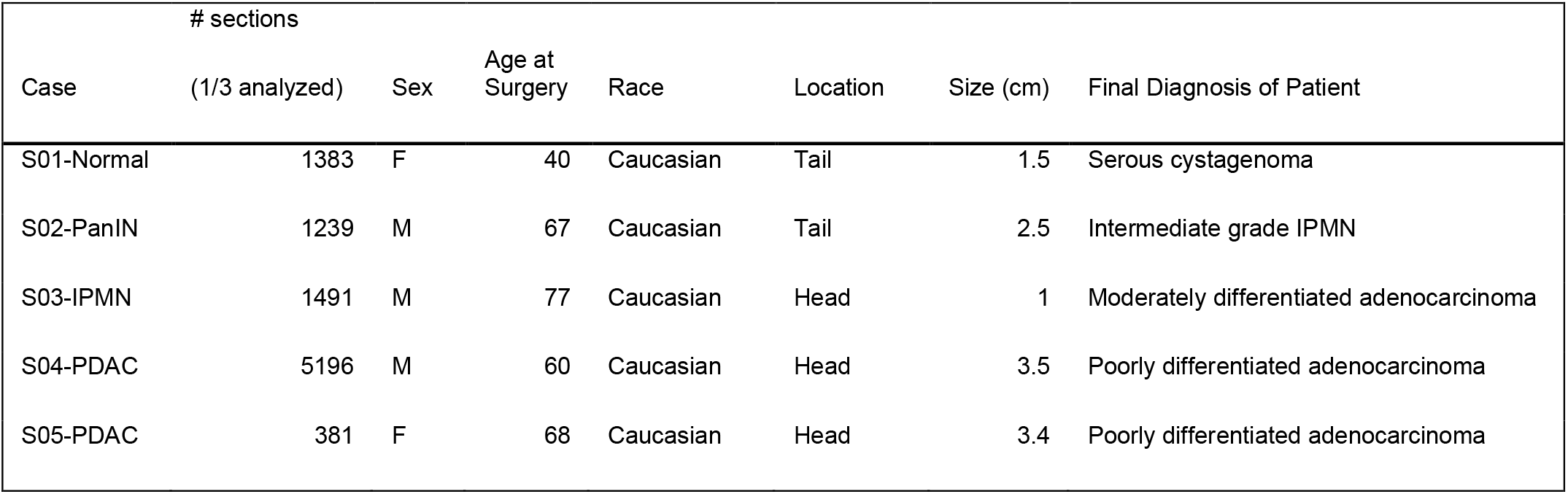
Patient Case information. Information about pancreas tissue samples analyzed. Tissues analyzed were adjacent normal, precancerous, or cancerous regions of human pancreas excised for clinical diagnosis.

## Supplementary Figures

**Supplementary Figure 1.**
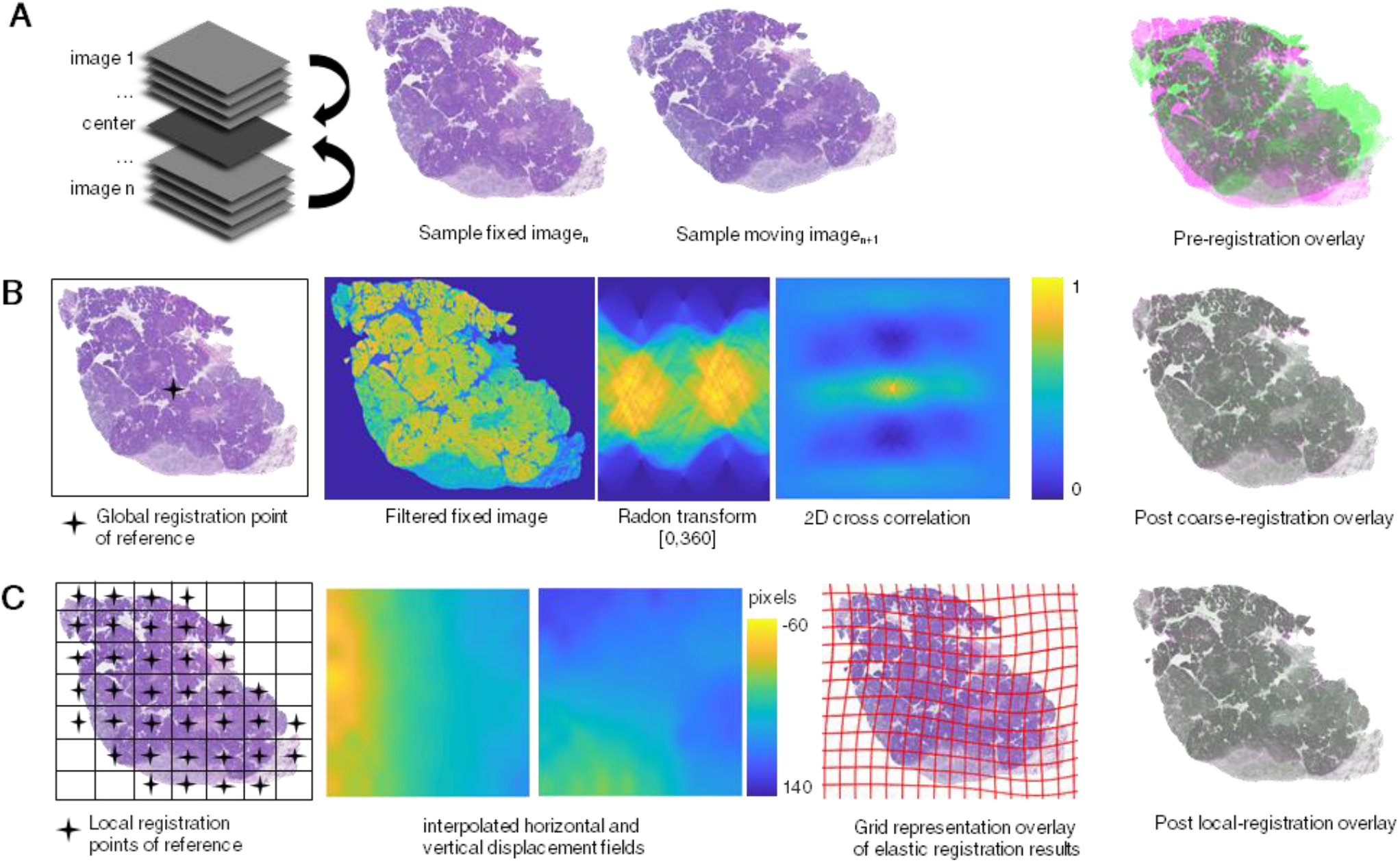
Histological image registration sample workflow. Sample images to illustrate image registration methodology. **A** Tissue cases registered with reference at center z-height of sample. Example fixed and reference image shown and overlaid. **B** Global registration performed with rotational reference at center of fixed image. Fixed and reference images smoothed by conversion to greyscale, removal of non-tissue objects in image, intensity complementing, and Gaussian filtering to reduce pixel-level noise in images. Radon transforms calculated filtered fixed and moving for discrete degrees 0-360. Maximum of 2D cross correlation of radon transforms yields registration angle. Maximum of 2D cross correlation of filtered images yields registration translation. Example global registered overlay. **C** Local registration performed at discrete intervals across fixed image. For each reference point, tiles are cropped from fixed and moving images and coarse registration is performed on tiles. Results from all tiles are interpolated on 2D grids to create nonlinear whole-image displacement fields. Overlay of fixed image and displacement grid exemplifies nonlinear registration results. Example local registered overlay.

**Supplementary Figure 2.**
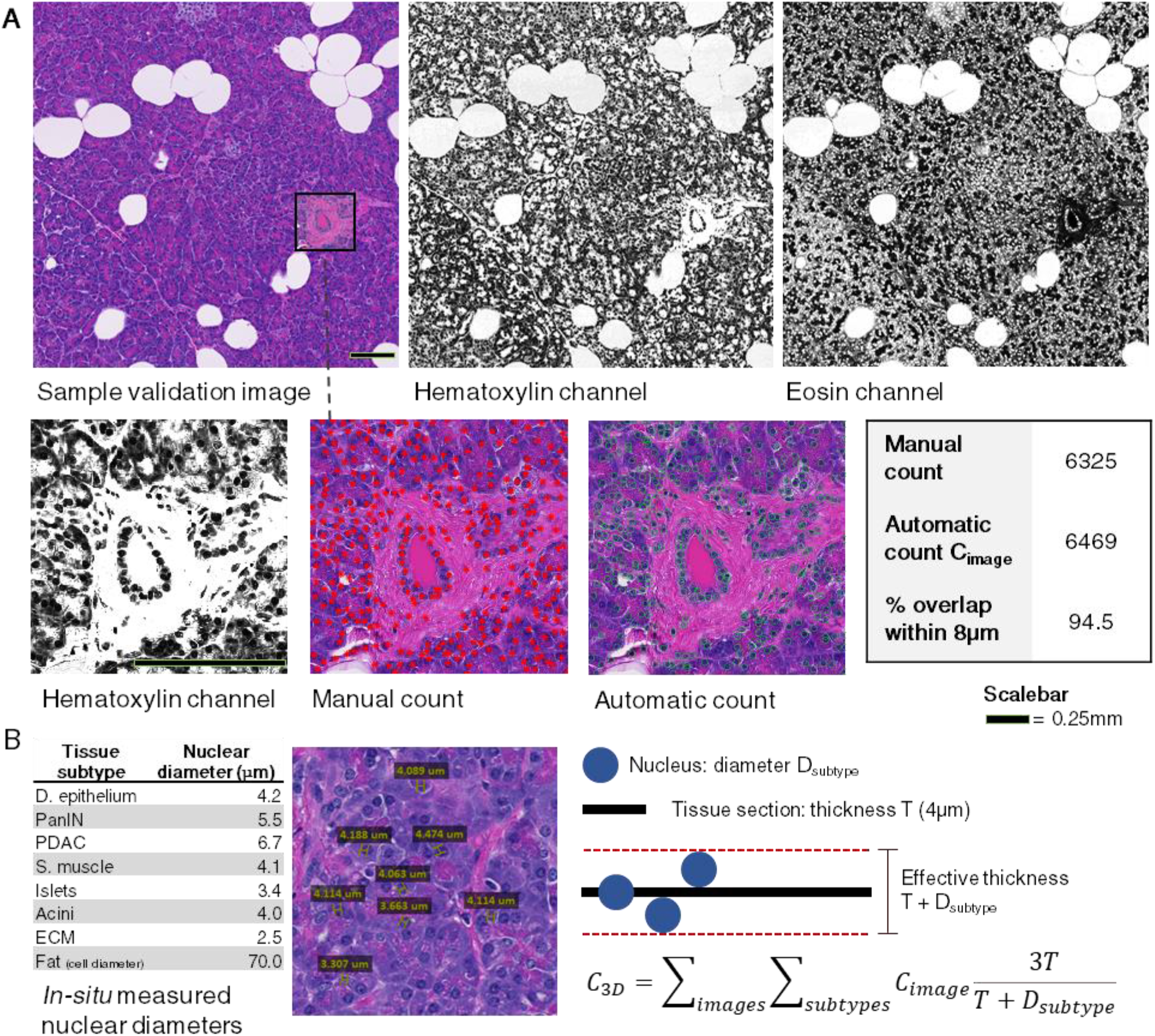
Validation of cell count and 2D to 3D cell count extrapolation. **A** Sample histological section and corresponding color deconvolved outputs representing hematoxylin and eosin channels of image. For 4mm^2^ tiles, calls were manually and automatically counted for validation of cell counting algorithm. 94% overlap was achieved between manual and automatic 2D cell counts **B** Cell diameters of each tissue subtype were measured using Aperio lmagaScope. 2D cell counts were extrapolated to 3D using the formula listed. It was assumed that calls could be detected by the algorithm if any part of the nucleus touched the tissue section. Therefore, affective tissue section thickness equals true tissue section thickness plus the diameter of the call. 3D cell counts were estimated by multiplying 2D cell counts by the true thickness of the tissue section, multiplied by three because two sections were skipped during scanning, divided by the affective thickness of the section.

**Supplementary Figure 3.**
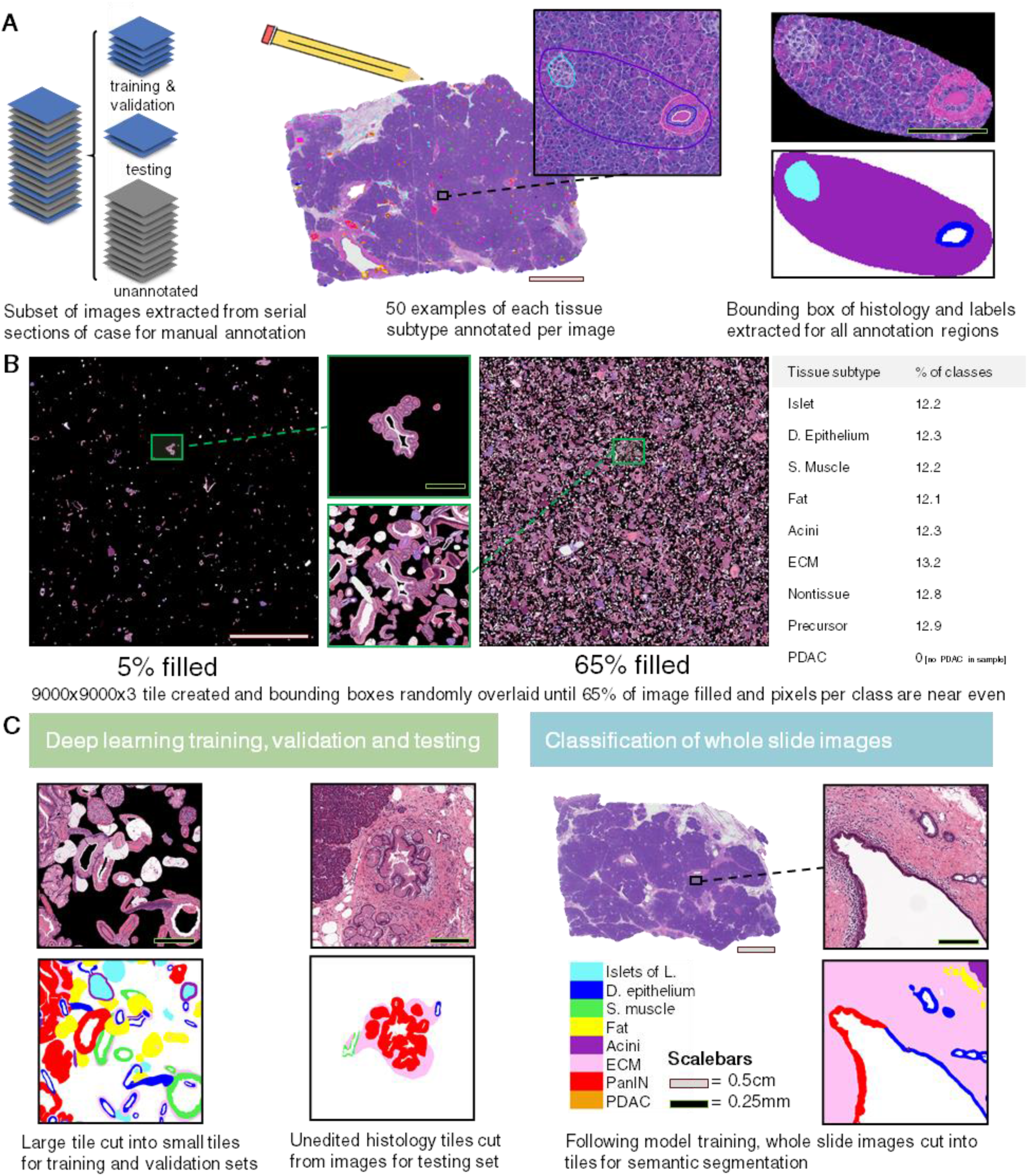
Overview of semantic segmentation workflow and training data design. **A** For each case, a minimum of seven images were extracted from for manual annotation. For each extracted image, minimum 50 examples of each tissue type were annotated, and the annotations cropped from the larger image. **B** To construct training and validation sets, cropped annotations were overlayed on a large image until the image was >65% full and such that the number of annotations of each type was roughly equal. **C** These large tiles were cut into smaller tiles for training and validation. Additional tiles were created for the testing set where the annotation was not cropped from the image. Testing accuracy was assessed as the percentage of the annotated area of the tile classified correctly. Following model training, the serial images were cropped into tiles and semantically segmented.

**Supplementary Figure 4.**
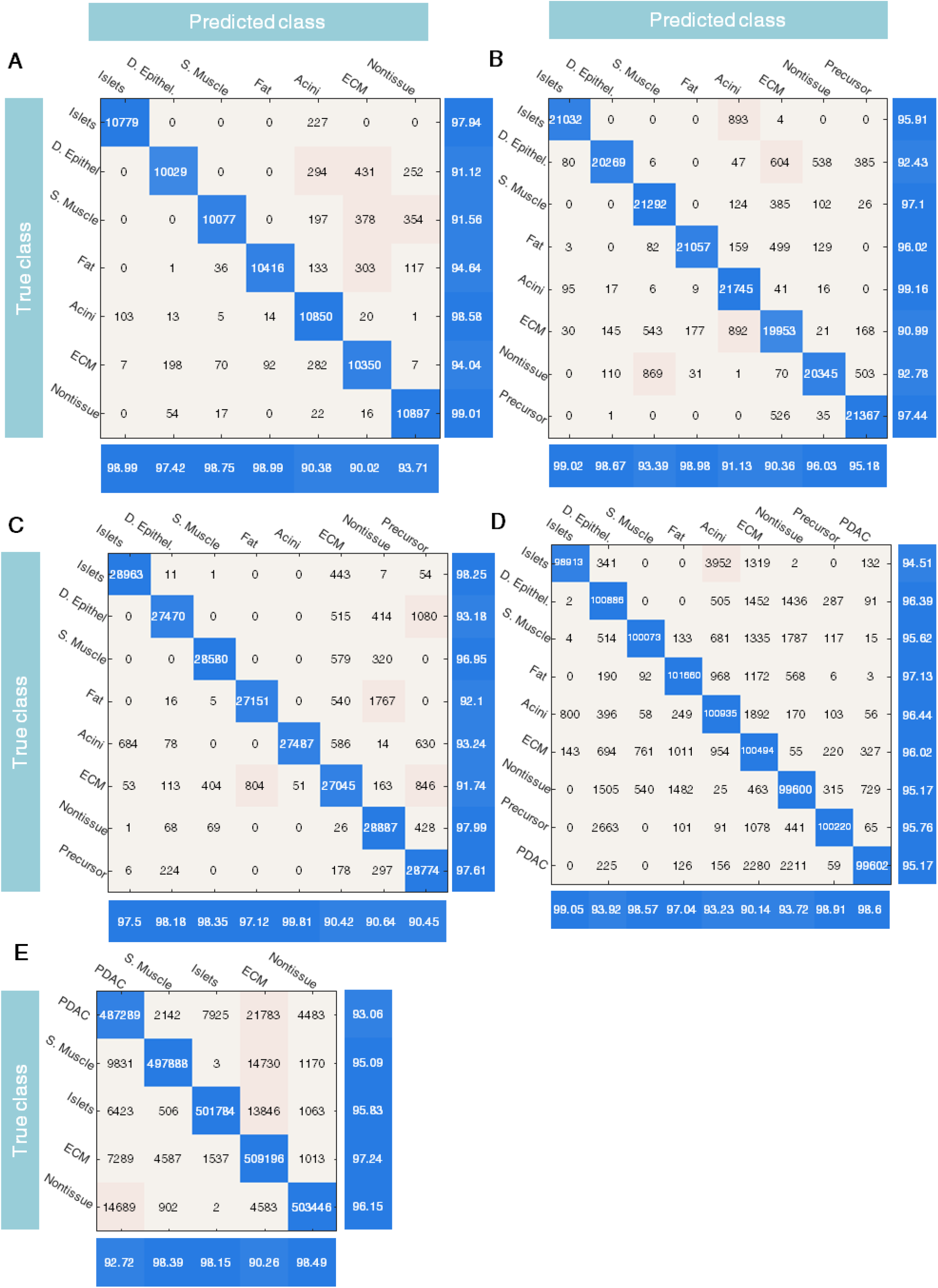
Deep learning accuracy for each tissue case. Confusion matrices display predicted vs. true (based on manual annotation) outcomes for all deep learning classes. Precision per class in row beneath matrices and recall per class in column to the right of matrices, both in units of percentage. **A** Results for S01-Normal. **B** Results for S02-PanIN. **C** Results for S03-IPMN. **D** Results for S04-PDAC. **E** Results for S05-PDAC.

## References

1. C. Liebig, G. Ayala, J. A. Wilks, D. H. Berger, D. Albo, Perineural invasion in cancer: a review of the literature. Cancer 115, 3379 (Aug 1, 2009).

2. S. M. Hong et al., Three-dimensional visualization of cleared human pancreas cancer reveals that sustained epithelial-to-mesenchymal transition is not required for venous invasion. Modern Pathol 33, 639 (Apr, 2020).

3. D. S. Michaud et al., Physical activity, obesity, height, and the risk of pancreatic cancer. Jama-J Am Med Assoc 286, 921 (Aug 22, 2001).

4. R. L. Siegel, K. D. Miller, A. Jemal, Cancer statistics, 2020. Ca-Cancer J Clin 70, 7 (Jan, 2020).

5. R. H. Hruban et al., Why is pancreatic cancer so deadly? The pathologist’s view. J Pathol 248, 131 (Jun, 2019).

6. L. Huang et al., Ductal pancreatic cancer modeling and drug screening using human pluripotent stem cell and patient-derived tumor organoids. Cancer Res 76, (Dec, 2016).

7. J. Drost, H. Clevers, Organoids in cancer research. Nature reviews. Cancer 18, 407 (Jul, 2018).

8. K. Taniuchi et al., Overexpressed P-cadherin/CDH3 promotes motility of pancreatic cancer cells by interacting with p120ctn and activating Rho-family GTPases. Cancer Res 65, 3092 (Apr 15, 2005).

9. R. Plentz et al., Inhibition of gamma-secretase activity inhibits tumor progression in a mouse model of pancreatic ductal adenocarcinoma. Gastroenterology 136, 1741 (May, 2009).

10. Z. Cruz-Monserrate et al., Detection of pancreatic cancer tumours and precursor lesions by cathepsin E activity in mouse models. Gut 61, 1315 (Sep, 2012).

11. B. Yang et al., Single-Cell Phenotyping within Transparent Intact Tissue through Whole-Body Clearing. Cell 158, 945 (Aug 14, 2014).

12. T. C. Murakami et al., A three-dimensional single-cell-resolution whole-brain atlas using CUBIC-X expansion microscopy and tissue clearing. Nat Neurosci 21, 625 (Apr, 2018).

13. E. A. Susaki et al., Versatile whole-organ/body staining and imaging based on electrolyte-gel properties of biological tissues. Nature communications 11, 1982 (Apr 27, 2020).

14. S. Zhao et al., Cellular and Molecular Probing of Intact Human Organs. Cell 180, 796 (Feb 20, 2020).

15. K. Chung et al., Structural and molecular interrogation of intact biological systems. Nature 497, 332 (May 16, 2013).

16. D. S. Richardson, J. W. Lichtman, Clarifying Tissue Clearing. Cell 162, 246 (Jul 16, 2015).

17. Y. Song, D. Treanor, A. J. Bulpitt, D. R. Magee, 3D reconstruction of multiple stained histology images. Journal of pathology informatics 4, S7 (2013).

18. J. M. Lotz et al., Integration of 3D multimodal imaging data of a head and neck cancer and advanced feature recognition. Biochimica et biophysica acta. Proteins and proteomics 1865, 946 (Jul, 2017).

19. J. Lotz et al., Zooming in: High Resolution 3D Reconstruction of Differently Stained Histological Whole Slide Images. Proc Spie 9041, (2014).

20. A. D. Singhi, E. J. Koay, S. T. Chari, A. Maitra, Early Detection of Pancreatic Cancer: Opportunities and Challenges. Gastroenterology 156, 2024 (May, 2019).

21. N. Roberts, Toward Routine Use of 3D Histopathology as a Research Tool (vol 180, pg 1835, 2012). Am J Pathol 181, 374 (Jul, 2012).

22. D. Magee et al., Histopathology in 3D: From three-dimensional reconstruction to multi-stain and multi-modal analysis. Journal of pathology informatics 6, 6 (2015).

23. P. H. Wu et al., High-throughput ballistic injection nanorheology to measure cell mechanics. Nature protocols 7, 155 (Jan 5, 2012).

24. J. Saltz et al., Spatial Organization and Molecular Correlation of Tumor-Infiltrating Lymphocytes Using Deep Learning on Pathology Images. Cell reports 23, 181 (Apr 3, 2018).

25. N. Coudray et al., Classification and mutation prediction from non-small cell lung cancer histopathology images using deep learning. Nature medicine 24, 1559 (Oct, 2018).

26. U. Djuric, G. Zadeh, K. Aldape, P. Diamandis, Precision histology: how deep learning is poised to revitalize histomorphology for personalized cancer care. Npj Precis Oncol 1, (Jun 19, 2017).

27. J. J. Nirschl et al., A deep-learning classifier identifies patients with clinical heart failure using whole-slide images of H&E tissue. Plos One 13, (Apr 3, 2018).

28. L.-C. Z. Chen, Yukun, Papandreou, George, Schroff, Florian Adam, Hartwig, Encoder-Decoder with Atrous Separable Convolution for Semantic Image Segmentation. Proceedings of the European Conference on Computer Vision (ECCV), pp. 801 (2018).

29. D. Hutchings et al., Cancerization of the Pancreatic Ducts: Demonstration of a Common and Under-recognized Process Using Immunolabeling of Paired Duct Lesions and Invasive Pancreatic Ductal Adenocarcinoma for p53 and Smad4 Expression. Am J Surg Pathol 42, 1556 (Nov, 2018).

30. K. Brune et al., Multifocal neoplastic precursor lesions associated with lobular atrophy of the pancreas in patients having a strong family history of pancreatic cancer. Am J Surg Pathol 30, 1067 (Sep, 2006).

31. R. H. Hruban et al., An illustrated consensus on the classification of pancreatic intraepithelial neoplasia and intraductal papillary mucinous neoplasms. Am J Surg Pathol 28, 977 (Aug, 2004).

32. C. R. Drifka et al., Periductal stromal collagen topology of pancreatic ductal adenocarcinoma differs from that of normal and chronic pancreatitis. Modern Pathol 28, 1470 (Nov, 2015).

33. L. Zhu, G. Shi, C. M. Schmidt, R. H. Hruban, S. F. Konieczny, Acinar cells contribute to the molecular heterogeneity of pancreatic intraepithelial neoplasia. Am J Pathol 171, 263 (Jul, 2007).

34. J. P. Morris, D. A. Cano, S. Selkine, S. C. Wang, M. Hebrok, beta-catenin blocks Kras-dependent reprogramming of acini into pancreatic cancer precursor lesions in mice. J Clin Invest 120, 508 (Feb, 2010).

35. S. S. Xu et al., The role of collagen in cancer: from bench to bedside. J Transl Med 17, (Sep 14, 2019).

36. T. J. Puls, X. H. Tan, C. F. Whittington, S. L. Voytik-Harbin, 3D collagen fibrillar microstructure guides pancreatic cancer cell phenotype and serves as a critical design parameter for phenotypic models of EMT. Plos One 12, (Nov 30, 2017).

37. C. R. Drifka et al., Highly aligned stromal collagen is a negative prognostic factor following pancreatic ductal adenocarcinoma resection. Oncotarget 7, 76197 (Nov 15, 2016).

38. S. I. Fraley et al., Three-dimensional matrix fiber alignment modulates cell migration and MT1-MMP utility by spatially and temporally directing protrusions. Scientific reports 5, 14580 (Oct 1, 2015).

39. S. P. Chiang, R. M. Cabrera, J. E. Segall, Tumor cell intravasation. American journal of physiology. Cell physiology 311, C1 (Jul 1, 2016).

40. A. C. Rios et al., Intraclonal Plasticity in Mammary Tumors Revealed through Large-Scale Single-Cell Resolution 3D Imaging. Cancer Cell 35, 618 (Apr 15, 2019).

41. M. F. Cuccarese et al., Heterogeneity of macrophage infiltration and therapeutic response in lung carcinoma revealed by 3D organ imaging. Nature communications 8, 14293 (Feb 8, 2017).

42. M. Orlich, F. Kiefer, A qualitative comparison of ten tissue clearing techniques. Histology and histopathology 33, 181 (Feb, 2018).

43. B. D. Lehmann et al., Refinement of Triple-Negative Breast Cancer Molecular Subtypes: Implications for Neoadjuvant Chemotherapy Selection. Plos One 11, e0157368 (2016).

44. R. A. Walker, Quantification of immunohistochemistry--issues concerning methods, utility and semiquantitative assessment I. Histopathology 49, 406 (Oct, 2006).

45. A. Goode, B. Gilbert, J. Harkes, D. Jukic, M. Satyanarayanan, OpenSlide: A vendor-neutral software foundation for digital pathology. Journal of pathology informatics 4, 27 (2013).

46. W. Falkena, xml2struct. (2020).

47. H. Hoffmann, Simple violin plot using matlab default kernel density estimation. INRES (University of Bonn), (2015).

